# Metabolic stability underpins thermal tolerance in benthic ecotypes of threespine stickleback (*Gasterostues aculeatus)*

**DOI:** 10.1101/2025.09.15.676414

**Authors:** Emily V. Kerns, Kristofer T. Sasser, Jesse N. Weber

## Abstract

Understanding how organisms adapt to divergent thermal conditions can inform efforts to preserve biodiversity in a warming world. Change in aerobic scope (AS)–total energy available to perform tasks outside of homeostasis–is an ecologically relevant predictor of performance across a range of temperatures. Mathematical models suggest that the shape of AS-temperature curves should evolve depending on environmental heterogeneity, but few studies have tested whether this occurs at the intraspecific level. We examined AS-temperature associations across two ecotypes of threespine stickleback fish (*Gasterosteus aculeatus*), including both short- and long-term heat treatments. We predicted that fish from shallow, thermally dynamic lakes would display greater AS stability at high temperatures than residents of deep, thermally stable lakes. We also tested whether heat and AS are correlated with variation in a fibrotic immune response after exposure to the cestode *Schistocephalus solidus*. We found few heat-associated differences in condition or immunity, but fibrosis decreased AS. Additionally, limnetic AS declined with heat and this shift was correlated with mortality. In contrast, benthic ecotypes displayed remarkable stability in AS and survival across temperatures. Connecting intraspecific variation in AS with local thermal environments is a promising avenue for estimating, and potentially fostering, population resilience to warming climates.

## Introduction

Global warming is altering species distributions (Chen et al., 2011) and driving population declines that can ultimately lead to extinctions (Hof et al., 2011; Sayer et al. 2025). These trends are particularly acute in species that inhabit freshwater environments (Sayer et al., 2025). Physiological plasticity is often proposed as a mechanism for resilience to global warming (Seebacher et al., 2015). However, theory regarding how plasticity may influence resilience to thermal challenge is conflicting. It is often argued that populations that live in variable environments are more likely to evolve adaptive plasticity, and therefore may be more robust to rapid environmental fluctuations (Botero et al., 2015). Mathematical models also predict high within generation shifts in temperature should select for more stable performance across a range of temperatures (Asbury & Angilletta, 2010), but the result requires tradeoffs between generalist versus specialist strategies that are not always supported by empirical data (Asbury & Angilletta, 2010; Angilletta et al., 2002). Alternatively, canalized phenotypes can sometimes be favored under fluctuating selection (Kawecki, 2000), and plastic populations may be more susceptible to heat stress (Angilletta et al., 2002; Bogan et al., 2024).

Although complex mechanisms underlie temperature-related shifts in metabolic rates (Clarke and Fraser, 2004; Schulte, 2015), researchers have tended to focus on measuring metabolic changes at thermal limits or performance curves over a range of temperatures. The oxygen- and capacity-limited thermal tolerance hypothesis (OCLTTH) integrates these approaches by focusing on physiological changes that occur up to critical thermal maximum (CT_max_), positing that thermal tolerance results from the plastic ability to increase transport of oxygen to tissues under elevated temperatures (Pörtner, 2001; Pörtner, 2002). The OCLTTH predicts that aerobic scope (AS), the difference between the minimum and maximum metabolic rate of an organism (Farrell, 2016; Fry, 1971; Fry, 1947), will increase until an optimal temperature (T_opt_) is reached, after which AS will decline (Supplemental Fig. 1A). The lower AS observed at temperatures above T_opt_ should result from more oxygen being necessary to maintain homeostasis (i.e. higher standard metabolic rate (SMR)) accompanied by either a stable or decreased maximum metabolic rate (MMR). Therefore, decreasing AS at higher temperatures should be among the first indicators of decreased performance.

Proponents of the OCLTTH emphasize that focusing on somatic maintenance provides clear ecological relevance (Pörtner et al., 2017). However, the OCLTTH remains contentious among ecophysiologists (Jutfelt et al., 2018). A meta-analysis of 73 studies found that almost half failed to find a thermal optima for AS, undercutting a major pillar of the hypothesis (Lefevre, 2016). This lack of support may stem from multiple causes, including natural variation in time-dependent physiological responses (e.g., the ability of organisms to acclimate to high temperatures (Fangue et al, 2006)). More specifically, when organisms encounter thermal stress they display variation in their primary (immediate), secondary (long term maintenance of homeostasis), and tertiary (dysregulation of biological systems) responses (Franke et al., 2024; Islam et al., 2022). Given that organismal fitness depends on the fluctuations of environments experienced throughout a lifetime, and that populations evolve in different environments, to understand how plasticity influences resilience requires testing physiological responses to stress in both temporal and population-specific contexts.

While hypotheses connecting respiration and temperature have received substantial experimental attention, less is known about the effect warming will have on other energetically demanding tasks such as immunity to parasite infection (Franke et al., 2024). One hypothesis is that global warming will exacerbate the impacts of parasitism by disrupting host immunity and facilitating parasite transmission, infectivity, and growth (Lõhmus & Björklund, 2015; Macnab & Barber, 2012). The combination of heat and parasite infection may exacerbate the detrimental effects of each stressor, synergistically amounting to a more negative physiological response for hosts (Franke et al., 2017; Franke et al., 2024). Alternatively, heat may detrimentally impact fitness of parasites (Kirk et al., 2022), hindering their ability to complete their life cycle (Claar & Wood, 2020; Barber et al., 2016).

Threespine stickleback (*Gasterosteus aculeatus*) fish provide an opportunity to test whether populations that evolved in different thermal habitats display metabolic differences related to thermal tolerance, immunity, and their interaction. Multiple studies have found heritable differences in thermal tolerance of threespine stickleback ecotypes (Barrett et al., 2011; Dittmar et al., 2014). Freshwater stickleback experience colder winter temperatures than marine fish and repeatedly evolved lower critical thermal minima (CT_min_: Barrett et al., 2011), but there is no evidence of evolved differences in CT_max_ (Barrett et al., 2011; Mottola et al., 2022). There is also substantial temperature variation across freshwater habitats. Benthic ecotypes are so named because they evolved to feed on benthic invertebrates that are plentiful in shallow lakes. In contrast, limnetic ecotypes are adapted to feed on zooplankton that are common prey in large, deep lakes (Willacker et al., 2010). These two environments should also differ in temperature: shallow lakes can warm and cool rapidly while deep lakes are more thermally stable. Given that many benthic and limnetic stickleback populations have inhabited divergent temperature environments for thousands of years, they offer an opportunity to test for the evolution of divergence in thermal tolerance. To our knowledge, only one study has tested whether thermal tolerance differs among stickleback sampled from lakes of different size and mean summer temperature (Dammark et al., 2018). This work found only slight differences in short term CT_max_ and did not explicitly categorize fish by morphological ecotype.

In addition to inhabiting thermally divergent environments, benthic and limnetic stickleback differ in parasite exposure. Limnetic ecotypes tend to consume more plankton than benthic ecotypes, which drives higher exposure to the copepod-transmitted cestode *Schistocephalus solidus* (Bolnick et al., 2026; Stutz et al., 2014). To minimize the impacts of *S.solidus* infection, many stickleback populations evolved to initiate, upon parasite exposure, an intense innate immune response that results in extensive peritoneal fibrosis (Hund et al. 2022; Weber et al., 2017b). However, severe fibrosis can be costly in terms of energy required to mount the response (Sasser et al., in review), likely contributing to the evolved loss of fibrosis in some populations (Weber et al., 2022). Conversely, the protective effects of fibrosis may be particularly beneficial under warming conditions. Higher temperatures increase both the growth rate and fecundity of *S. solidus* (Franke et al., 2017; Macnab & Barber, 2012; Scharsack et al., 2021). One caveat is that some stickleback populations exhibit immune dysregulation at hotter temperatures (Scharsack et al., 2021), even in the absence of pathogens (Dittmar et al., 2014). Though complicated, these immune-temperature connections should be most apparent in limnetic stickleback ecotypes, as they are expected to display both high cestode immunity (i.e., strong fibrosis) and limited thermal resilience.

We conducted an experiment to test three primary questions (Supplemental Fig. 1A): 1) Do stickleback metabolic rates conform to the predictions of the OCLTTH, 2) do benthic and limnetic stickleback differ in their primary, secondary, and tertiary responses to heat, and 3) do increased temperatures impair fibrotic immune responses, thereby affecting parasite infections, AS, and/or fish condition? According to the OCLTTH, we expected AS to reach an optimal level and then decline at higher temperatures due to increased SMR without a corresponding decrease in MMR. This decrease in AS should be accompanied by declines in condition metrics. We predicted that benthic ecotypes would show greater thermal tolerance, reflected by less change in AS with temperature. This could be accomplished either through smaller increases in SMR or an ability to maintain high MMR across a broader range of temperatures. While we initially defined ecotypes based on morphological characters, in reality populations display a continuum of traits connected to environmental differences (Haines et al., 2022). Indeed, recent work suggests that fibrosis is best predicted by physical lake characteristics (Bolnick et al., 2026). Therefore, we also tested whether lake properties (i.e., depth and surface area) predicted variation in thermal tolerance. With respect to immunity, we expected that higher temperatures would weaken fibrosis responses and increase the incidence of cestode infections due to higher parasite growth rates at warmer temperatures. When fibrosis did occur, we expected it to drive reduced AS and body condition. Finally, we expected heat-induced decreases in fibrosis to be most pronounced in cestode-immune limnetic fish.

## Methods

### Animal husbandry and thermal manipulations

We generated experimental fish via *in vitro* fertilization of gametes from wild-caught parents. The parents were sampled from four lakes in southcentral Alaska (Supplemental Table 1): Finger and Watson lakes are shallow (13.4m and 4.3m, respectively), while Spirit and Wik are deep (21m and 24.4m, respectively). Morphological analyses assigned Watson and Finger as benthic ecotypes, Spirit as moderately limnetic, and Wik as highly limnetic (Haines et al., 2022; Hendry et al., 2024). Before the experiment fish were maintained at a temperature of 17 ± 1°C.

During the experiment, 16-19-month old adult fish were placed into mixed-family, population-specific tanks. Each population was represented by 3-4 families that were spread evenly across temperature treatments. We acclimated the fish to their experimental treatments (18-26℃) by increasing water temperature 1℃ per day (Supplemental Fig. 1B). After acclimation, we maintained fish at experimental temperatures for 31-32 days (Supplemental Fig. 1B; details below). Fish were monitored daily, and any animals showing signs of distress (i.e., erratic swimming, immobility, loss of equilibrium, or gasping) were euthanized using an overdose of pH buffered MS-222. We similarly euthanized all fish at the end of the experiment and performed dissections to measure infection outcome. Sample sizes for all treatments are reported in Supplemental Table 1. Field sampling and lab experiments were approved by the Alaska Department of Fish and Game (Permit SF2022-043d) and University of Wisconsin-Madison IACUC (protocol #L006460).

### Respirometry

We measured mass specific SMR and MMR via intermittent flow respirometry (Supplementary methods) to assess primary (1-2 days post-acclimation) and secondary (31-32 days post-acclimation) physiological responses to heat. R:FishResp (Svendsen et al., 2019) was used to calculate AS as the difference between SMR and MMR (Clark et al. 2013). Respirometry traits were only measured in Watson (benthic) and Spirit (limnetic) populations at the secondary timepoint. Trials with equipment failures or mortality were excluded from later analyses.

### Experimental cestode exposures

After initial respirometry assays, we transferred individual fish into temperature-controlled containers and exposed them to infected copepods containing a total of ∼7 cestodes. Following Weber et al. (2022), batches of cyclopoid copepods (*Acanthocyclops robustus*) were exposed to lab-reared *S. solidus* coracidia descended from parasites caught in Walby Lake, Alaska (61.6178, −149.2138). We measured copepod infection rates by screening a subset of each batch two weeks post exposure. At the time of exposure, the fish had been fasted for 48 hours. To guarantee the opportunity to observe uninfected fish, 86% of the fish were exposed to parasites. We could not distinguish exposed from unexposed individuals in the final analysis. If a fish died or was euthanized prior to the end of the experiment, it was dissected to assess *S. solidus* presence.

### Health indices, infection outcome, and fibrosis

Spleenosomatic index (SSI - a metric of immune condition), hepatosomatic index (HSI - a proxy for energy storage), and gonadosomatic index (GSI - for males only, informative for reproductive investment) represent the percent of an organism’s total wet mass contained within the relevant organ (i.e., organ mass/wet mass * 100). We estimated body condition (i.e., fat relative to body length) using Fulton’s Index *K* (Bolger and Connolly, 1989), where *K* = (mass (mg)/standard length (mm)^3^) x 100,000, and scaled mass index (SMI; Pieg & Green, 2009). During dissections we screened for cestode infection and scored fibrosis following Hund et al. (2022). Ordinal scores were 0 for no visible fibrosis, 1 if organs were attached to the swim bladder, 2 if organs were fused together, 3 if organs were fused to the side of the body cavity, and 4 if the body cavity was entirely filled with fibrotic tissue.

### Statistical Analysis

We used generalized linear models to test whether primary and secondary (i.e., start and end of experiment) respirometry traits (MMR, SMR, and AS) responded to temperature, ecotype, and temperature*ecotype interactions. For secondary trials we were also able to test whether metabolic rates were influenced by fibrosis (presence and score) and sex. We fit linear mixed effect models (R package nlme v. 31-164; Bates et al., 2015) to test whether condition metrics (Fulton’s K, SMI, SSI, GSI, HSI) were predicted by AS, ecotype, and their interaction. Sex was included as a fixed effect and tank density (number of fish per tank summed over the duration of the experiment) as a random effect. In the absence of evidence to support the OCLTTH (i.e., no effect of AS), temperature could still influence condition metrics through other biological pathways. Therefore, we again used linear mixed effect models to test the effects of temperature, ecotype, temperature*ecotype, and sex on each condition metric, with tank density as a random effect. Cox regression was used to assess whether survival varied between ecotypes, across temperature treatments, or between ecotypes within a temperature treatment. Additional details regarding statistical analyses, respirometry assays, and husbandry can be found in the supplementary information.

## Results

### Primary Metabolic Response: Ecotype and lake ecology interact with temperature to explain variation in respirometry rates

Immediately following acclimation, MMR was best explained by a linear increase with temperature (Fig. 1A, Supplemental Table 2 & 3). MMR also varied depending on the surface area of source lakes (Supplemental Table 2 & 4, Supplemental Fig. 2A). Specifically, MMR increased more with temperature in populations that originated from lakes with a smaller surface area (temperature * surface area: t = −2.023, p = 0.0445). Quadratic temperature had the strongest effect on SMR (t = 3.494, p = 0.0006), but linear temperature also significantly affected SMR (t = −2.636, p = 0.00913) (Fig. 1B, Supplemental Table 2 & 3). Mean lake depth, not surface area, best predicted temperature-related shifts in SMR, but had no effect (Supplemental Table 2 & 4, Supplemental Fig. 2B). Although ecotype was not a significant predictor of either primary MMR or SMR, there were several notable patterns at higher temperatures. MMR of limnetic fish plateaued at 24℃ but continued to increase at 26 ℃ in benthic fish. In contrast, SMR increased 84.7% (619 to 1143 mgO_2_/kg/hr) between 22℃ and 24℃ in limnetic fish, but only increased 24.9% (733 to 916 mgO_2_/kg/hr) in benthic fish.

**Figure 1:**
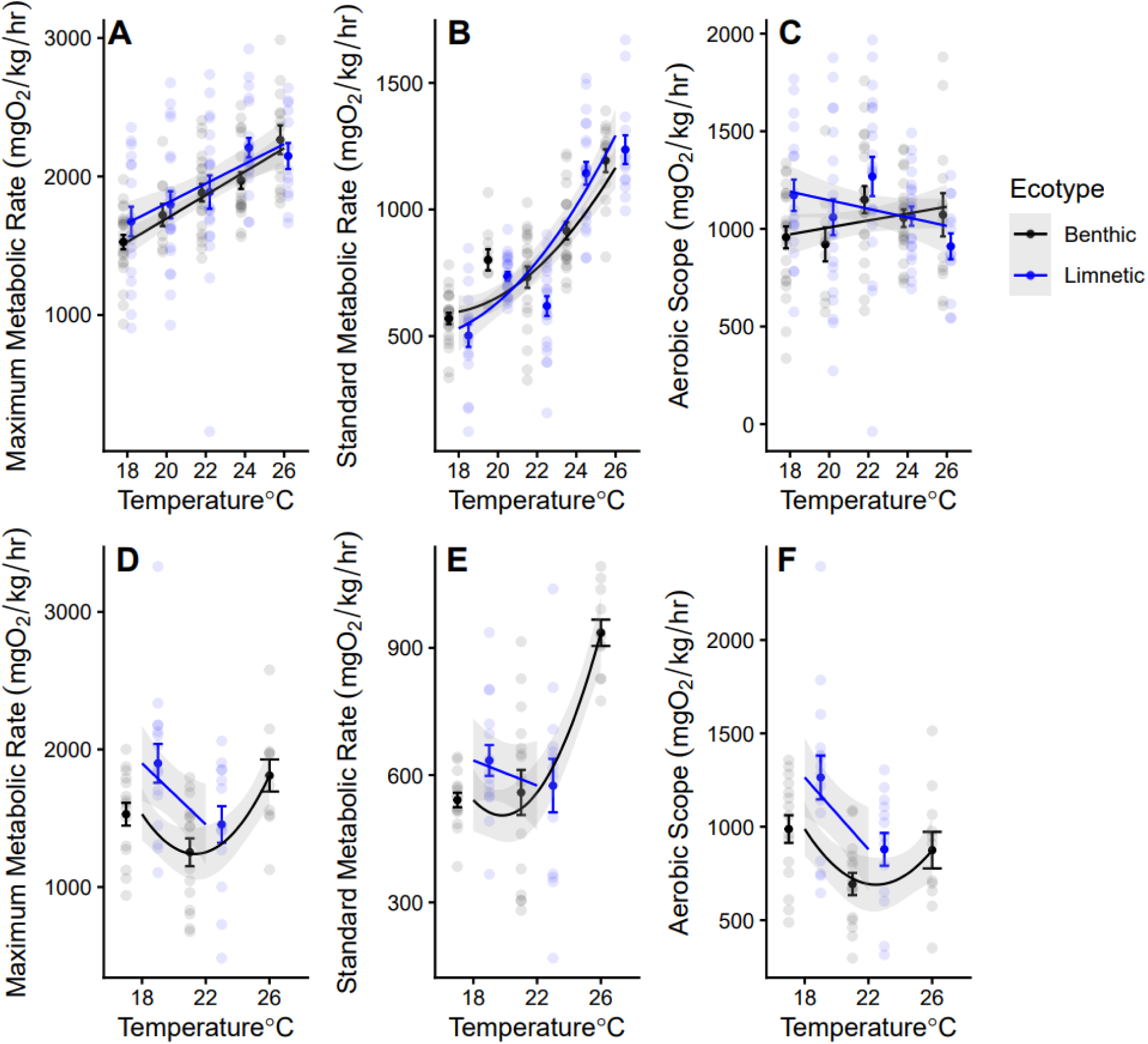
Metabolic data from the start **(A-C)** and end **(D-F)** of the experiment. **(A)** MMR and **(B)** SMR increased with temperature. **(C)** Limnetic fish displayed an overall higher AS, which increased to a peak at 22℃ followed by a sharp decline. The initial AS of benthic fish increased with temperature. After 31-32 days, **(D)** MMR, **(E)** SMR, and **(F)** AS decreased from 18℃ to 22℃ in both ecotypes, but then increased at 26℃ for benthic fish.

We next integrated MMR and SMR to estimate AS at each temperature. Although AS appeared to peak at 22°C in both ecotypes, only ecotype and temperature * ecotype effects were significant predictors in our model (Fig. 1C, Supplemental Table 2 & 3). When comparing AS between 18-22℃, limnetic fish had significantly higher AS than benthic fish (ecotype: t = 2.335, p = 0.0206), but above 22°C AS decreased precipitously in only limnetic fish (temperature * ecotype: t = −2.180, p = 0.0305). Indeed, limnetic fish had 18.3% higher AS at 18℃ than benthic fish, but 17.7% lower AS at 26℃. However, temperature * ecotype was an overall poor predictor of AS variation (R^2^_adj_ = 0.02, p = 0.086). The AS model with lake surface area, in place of ecotype, provided a better fit to our data (ΔAIC: 19.325, R^2^_adj_ = 0.128, p < 0.0001; Supplemental Table 2 & 4, Supplemental Fig. 2C); however, with only four populations we had limited ability to thoroughly test for connections between metabolism and continuous environmental variables.

### Secondary & Tertiary Response: Limnetic fish display substantial mortality at high temperature related to decreases in AS

Although this experiment was not designed to test survival under heat stress, mortality rates significantly increased with chronic exposure to higher temperature (z = 2.091, p = 0.0365), with the greatest effect on limnetic fish (temperature*ecotype: z = 1.949, p = 0.0512, Fig. 2). Much of this ecotype difference was driven by the 89% mortality of limnetic fish at 26℃, compared to just 11% mortality among benthic fish. Both benthic (t = −2.123, df = 94.165, p-value = 0.0364) and limnetic fish (t = −2.252, df = 45.628, p-value = 0.0292) showed significant declines in body condition between the start and end of the experiment (Supplemental Fig. 3). Across all temperatures, benthic fish experienced a 6.6% [95% CI: 0.42, 12.7%] decrease in body condition, while limnetic fish displayed a 7.3% [0.78, 13.9%] decline in body condition. Importantly, the limnetic decline in body condition is likely an underestimate given the paucity of individuals surviving at 26℃.

**Figure 2:**
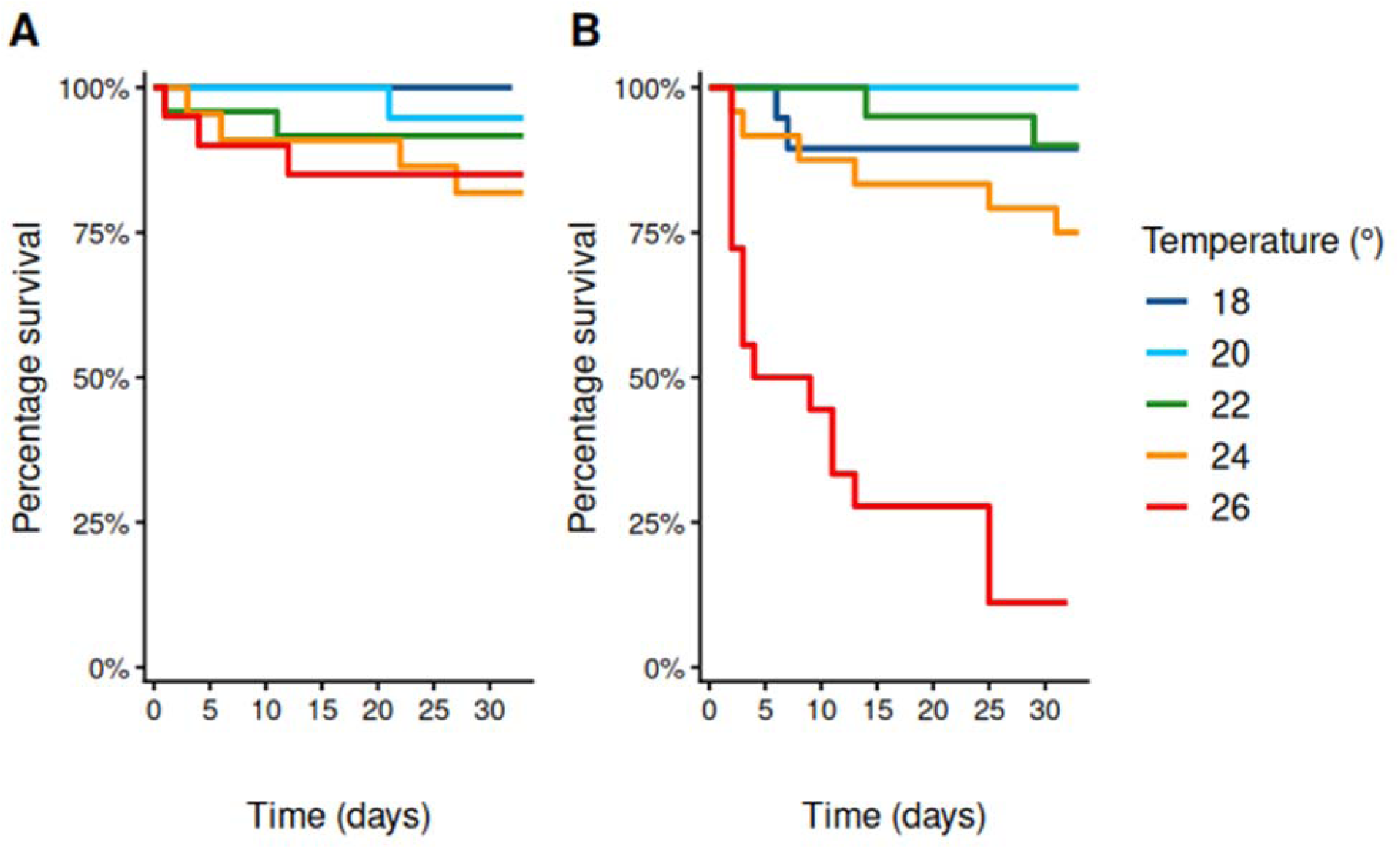
Probability of survival as function of temperature and time for **(A)** benthic and **(B)** limnetic stickleback. Survival probability significantly decreased at higher temperatures (z = 2.091, p = 0.0365) and there was a marginally significant effect of temperature*ecotype (z = 1.949, p = 0.0512).

Although we only examined secondary metabolic rates in two populations at three temperatures, we found significant linear and quadratic effects of temperature on MMR, SMR, and AS. Metabolic rates reached a minimum at 22℃ (Fig. 1D-F, Supplemental Table 2 & 5), with the only exception being mean SMR of benthic fish, which appeared to increase with heat. MMR (ecotype: t = 3.233, p = 0.00195) and AS (ecotype: t = 3.179, p = 0.002276) was significantly higher in limnetic fish than benthic fish at 18 and 22℃, but we could not compare rates at 26℃ due to limnetic mortality.

### Metabolic cost of fibrosis despite limited cestode infection across temperatures

Despite exposing 178 fish to *S. solidus*, we only recovered a single infected fish that died before the end of the experiment. We did, however, find that 72 fish exhibited peritoneal fibrosis, including individuals from all four populations. We also observed a few *S. solidus* encysted in fibrotic tissue. Although some stickleback populations, including Finger Lake, can sometimes display fibrosis without parasite exposure (Bolnick et al., 2026), observing fibrosis in 34% of cases suggests that exposures were successful but the parasites failed to establish infection after burrowing into the body cavity of the fish. Contrary to our prediction, temperature had no effect on fibrosis presence or severity for either ecotype (Supplemental Fig. 5D-E). However, as predicted, limnetic fish tended to fibrose more often than benthic fish, although this effect was only marginally significant (OR: 1.71, 95% CI [0.93, 3.18], p = 0.0877). Additionally, when either fibrosis presence or severity were included as predictors in MMR, SMR, and AS models (Supplemental Table 2), they had significant effects on all respirometry traits. Fibrosis presence was associated with lower MMR (t = −2.993, p = 0.0039), SMR (t = −2.952, p = 0.0044), and AS (t = −2.369, p = 0.0209), regardless of fish ecotype (Fig. 3). Fibrosis severity also led to a significant decline in MMR (t = −2.278, p = 0.0256) and SMR (t = −2.761, p = 0.0071), but not AS across both ecotypes (Supplemental Fig. 5A-C).

**Figure 3:**
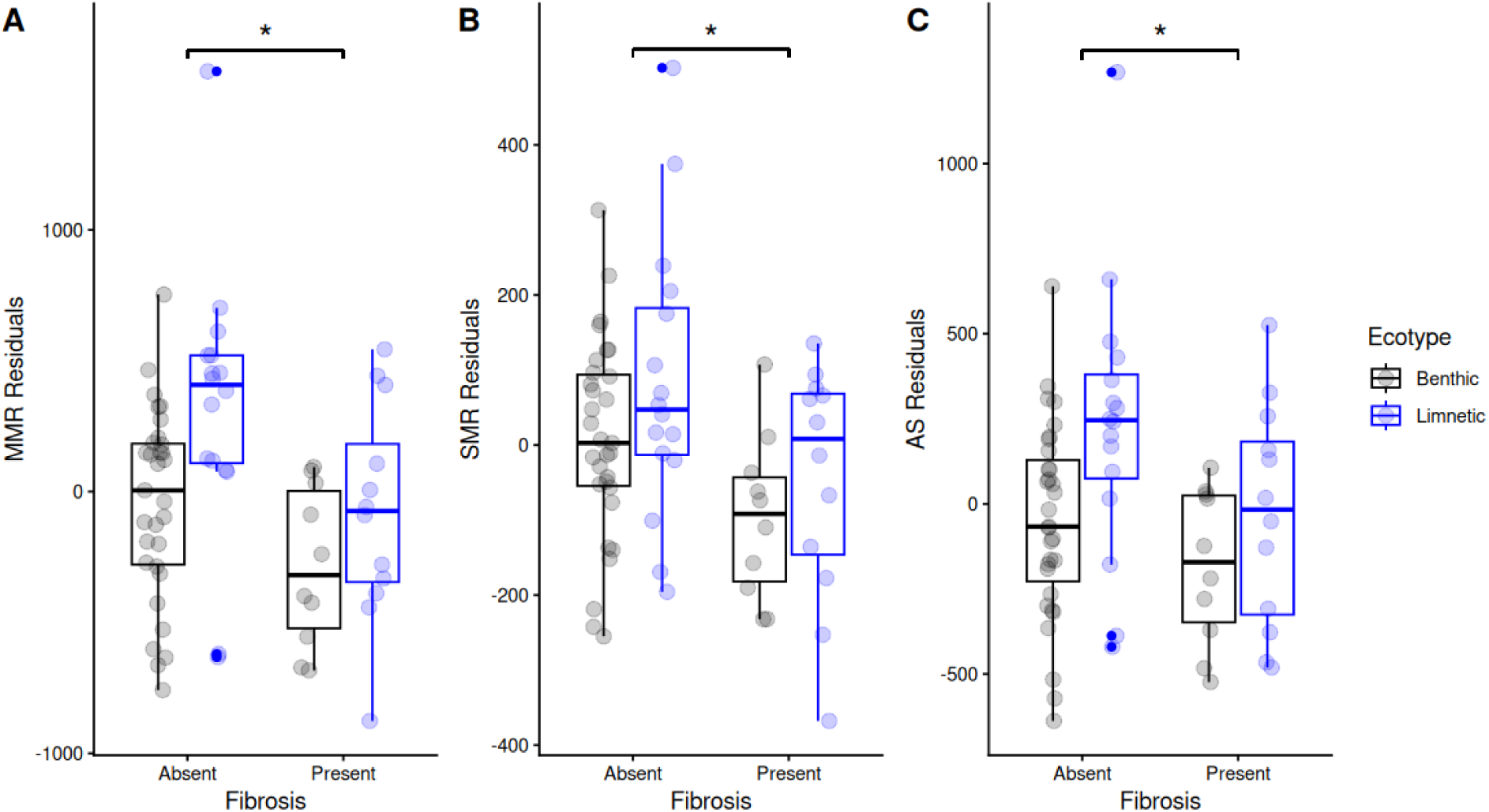
Residuals from the models for MMR, SMR, and AS with the effect of temperature and sex removed to display variation due to fibrosis and ecotype. **(A)** MMR, **(B)** SMR, and **(C)** AS were significantly lower in fish with peritoneal fibrosis, averaged across ecotypes. *p<0.05

### AS and temperature have ecotype- and population-dependent effects on some condition metrics

Based on the OCLTTH, we predicted that declines in AS would lead to reduced somatic condition. At both the primary (Fig. 4A) and secondary timepoint (Fig. 4B), the relationship between Fulton’s K body condition and AS depended on ecotype. Post-acclimation, limnetic fish showed an increase in body condition with AS (slope = 5.80 x 10^−5^ [95% CI: −5.51 x 10^−5^, 1.71 x 10^−4^]) while condition decreased in benthic fish (slope = −1.33^−4^ [−2.81 x 10^−4^, 1.47 x 10^−5^]; AS * ecotype: t = −2.026, df = 179, p = 0.0442). This trend reversed at the end of the experiment (AS * ecotype: t = 2.254, df = 61, p = 0.0278); Fulton’s K body condition increased with AS in benthic fish (slope = 1.12 x 10^−4^ [−4.71 x 10^−5^, 2.72 x 10^−4^]) but decreased in limnetic fish (slope = −1.19 x 10^−4^ [−2.50 x 10^−4^, 1.12 x 10^−5^]). The relationships between SMI and AS were identical to those obtained using Fulton’s K (Supplemental Fig. 8A-B).

**Figure 4:**
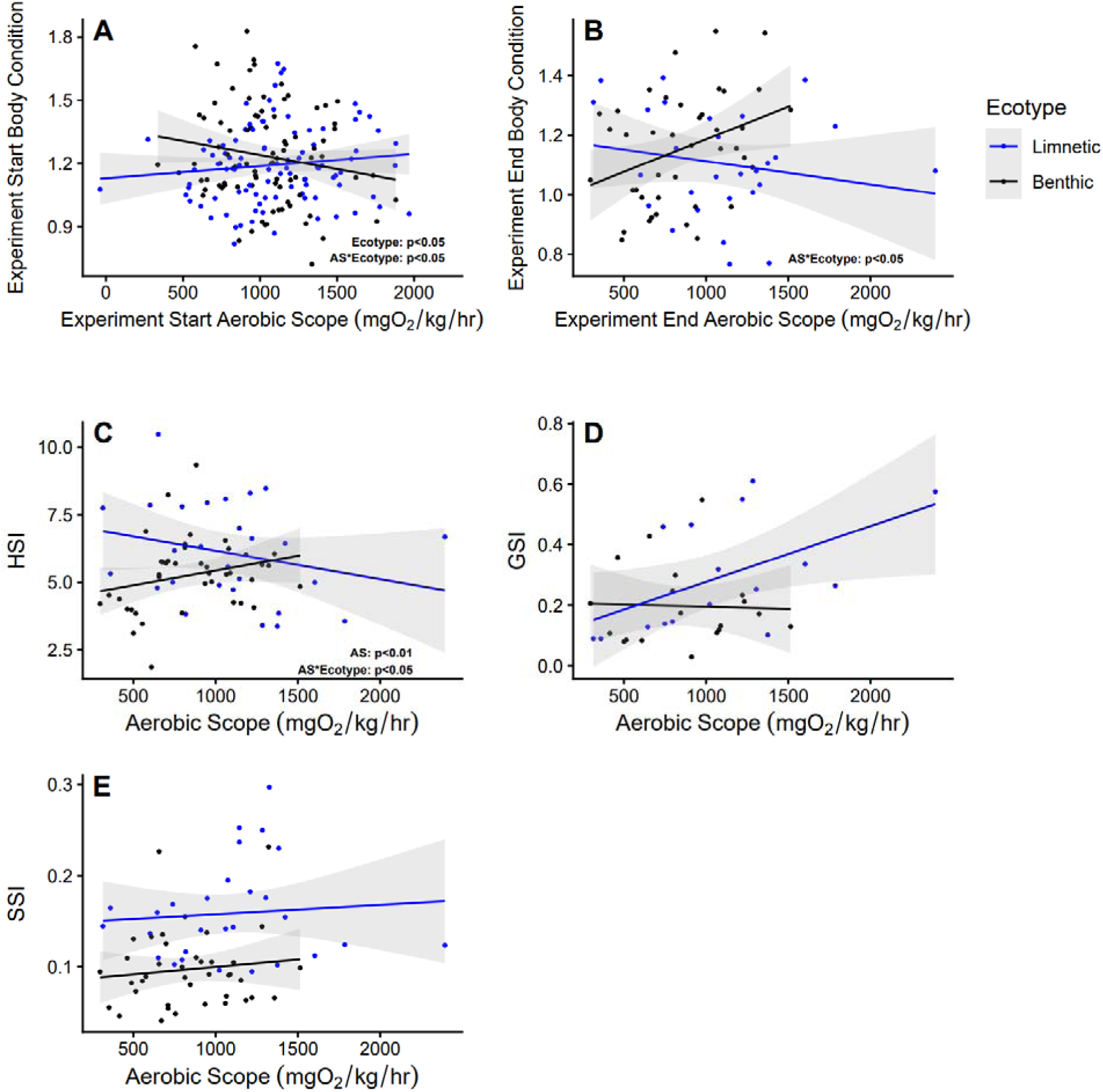
Individual condition metrics as a function of aerobic scope. Fulton’s K body condition at the primary **(A)** and secondary **(B)** timepoint showed opposite but significant ecotype-specific trends with AS. **(C)** HSI increased with AS for benthic fish but decreased in limnetic fish. Limnetic fish displayed a nonsignificant increase in GSI **(D)** while SSI **(E)** was stable for both ecotypes across AS.

As sex and all other condition metrics required lethal sampling, we only collected these data at the end of the experiment. Males had a higher body condition than females regardless of AS (t = 5.972, df = 61, p < 0.0001, Supplemental Fig. 6). Like body condition, HSI increased with AS in benthic fish (slope = 2.114 x 10^−3^ [5.43 x 10^−4^, 3.68 x 10^−3^]) but not limnetic fish (slope = −2.42 x 10^−4^, [−1.524 x 10^−3^, 1.04 x 10^−3^]; AS * ecotype: t = 2.322, df = 60, p = 0.0236, Fig. 4C). Unlike body condition, females had a higher HSI than males regardless of AS (t = −2.9996, df = 60, p = 0.0039, Supplemental Fig. 6B). GSI and SSI (Fig. 4D-E) had no relationship with AS for either ecotype.

Lastly, we assessed if temperature shifts had a direct effect on condition. Although separate from the OCLTTH, we could utilize data from all four populations to test these associations. SMI decreased significantly more in limnetic fish than benthic fish at the start of the experiment (temperature * ecotype: t = 2.084, df = 192, p = 0.0385, Supplemental Fig. 8C). However, there were no significant effects of temperature on Fulton’s K at the primary timepoint (Supplemental Fig. 4A). At the secondary timepoint, Fulton’s K (temperature * ecotype: t = −2.108, df = 152, p = 0.0366, Supplemental Fig. 4B) and SMI (temperature * ecotype: t = −2.228, df = 152, p = 0.0273, Supplemental Fig. 8D) decreased slightly more at higher temperatures in benthic fish than limnetic fish, but both were overall higher in benthic fish (Fulton’s K ecotype: t = 2.417, df = 152, p = 0.0168; SMI ecotype: t = 2.463, df = 152, p = 0.0149). There were no direct effects of temperature on HSI, SSI, or GSI (Supplemental Fig. 4C-D). When assessed at the population-level (Supplemental Fig. 7), only Finger Lake fish had a significant association between Fulton’s K body condition and heat (slope = 0.04648 [0.0218, 0.0712]; Supplemental Fig. 7A). However, by the end of the experiment there was no association between body condition, HSI, GSI, and SSI with temperature regardless of population (Supplemental Fig. 7B-E).

## Discussion

Thermal tolerance is used to project changes in geographic range (Sunday et al., 2012; Smith et al., 2022), population declines (Roeder et al., 2021), and identify species at risk of either local or global extinction (Turko et al., 2021). However, studies often assess tolerance in relation to critical thermal maxima (CT_max_), which is usually assayed by steadily increasing temperature until an organism is unable to maintain equilibrium. While this approach identifies a short-term thermal limit, it can obscure more ecologically-relevant differences caused by prolonged exposure to less extreme temperatures. Additionally, thermal tolerance may be affected by additional stressors that wild individuals are likely to experience simultaneously, such as pathogen exposure. Our experiment addressed these issues by examining habitat-associated metabolic responses to not only acute and chronic thermal stress, but also to parasite exposure and resulting immune responses. Consistent with our ecological prediction, we found that benthic populations were indeed more resilient to hotter temperatures, likely resulting from their ability to maintain stable AS. However, we found mixed evidence for the somatic decline connected to reduced AS, as expected by the OCLTTH. The association between AS and body condition was both ecotype and temporally-variable, showing opposite trends between the primary and secondary timepoint. Although we observed no successful cestode infections nor any impact of temperature on fibrosis, MMR, SMR, and AS decreased in fibrotic fish. This suggests that immune responses themselves may play an underappreciated role in shaping natural variation in metabolic capacity and thermal tolerance.

### Stable AS may enable benthic populations to survive at high temperatures

As noted previously, we designed our temperature challenge to fall well below the reported CT_max_ for stickleback. Another study that tested for associations between natural thermal environment and heat tolerance across populations of freshwater stickleback found no clear evidence for local adaptation and a general CT_max_ of 31-32°C (Dammark et al., 2018). This same experiment also found only modest transcriptomic evidence for heat stress in fish housed at 26°C for up to 48 hours. In contrast, in our chronic thermal challenge we observed significantly lower survival at higher temperatures, with the impact being most severe in the 26°C limnetic treatment (Fig. 2). It is unlikely that wild Alaskan stickleback will be continuously exposed to this high of temperature, and all populations showed high survival within a few days of acclimation. Regardless, our results highlight that stickleback populations respond differently to prolonged temperature increases, with limnetic fish suffering potentially large and negative consequences from even relatively small (i.e., compared to CT_max_) but prolonged temperature shifts. These longer term consequences were related to ecotype-specific metabolism differences observed immediately after acclimation. At the start of the experiment, AS peaked in limnetic fish at 22°C and declined by 28% at the highest temperature. Another study on long-term thermal responses used a different limnetic population from Alaska and, though not spanning the same temperatures, also found substantial plasticity in AS (Ressel et al., 2021). Meanwhile, initial AS remained relatively stable across all experimental temperatures for benthic fish, declining by only 6.7% between 22°C and 26°C. It is important to note that the trajectory of change in AS with temperature was more telling than absolute AS values. Both ecotypes had equivalent measures of AS at 24°C, but the downward trajectory of limnetic AS likely reflects the early stages of physiological dysregulation (i.e. tertiary response).

Although our chronic heat metabolic data are limited to two populations and three temperatures, they contain several interesting trends. The secondary benthic AS-temperature curve shifts to a convex shape, but likely the more important relationship is the change between primary and secondary AS values (Fig. 1). The difference in AS at 18℃ was negligible for both populations, but at 22℃ AS decreased by ∼47% and ∼22% in benthic Watson Lake fish and limnetic Spirit Lake fish, respectively. This finding suggests that Watson fish were beginning to show signs of stress following 30+ days of chronic heat exposure. Notably, we were unable to assay any limnetic fish at 26℃ at the secondary timepoint because this temperature was beyond their threshold of tolerance, highlighting that physiological responses result from the interaction between genetic background and length of time exposed to the stressor (Islam et al., 2022; Fangue et al. 2006; Franke et al., 2024). Altogether, immediate physiological dysregulation following acclimation likely led to higher mortality rates in limnetic populations while benthic fish were able to maintain homeostasis, enabling survival at higher temperatures.

### Ultimate and proximate causes of ecotype differences in thermal tolerance

How did the two ecotypes come to differ in their thermal tolerance? Given that all experimental fish were reared under common lab conditions and our results match ecological expectations for the two ecotypes, the most parsimonious explanation is that differences in thermal tolerance are heritable and likely arose via local adaptation of benthic fish to warmer water temperatures. Indeed, a previous study induced evolution of CT_min_ after just three years in marine stickleback (Barrett et al., 2011), highlighting the evolutionary potential of thermal performance curves.

Understanding the proximal causes of thermal tolerance differences will require disentangling variation in observed metabolic traits from variation in underlying cellular mechanisms. Although we observed notable stability of AS in benthic fish, this may have been facilitated either by greater plasticity in their ability to shunt energy reserves toward anaerobic metabolism (e.g., Hao et al., 2024) or loss of maladaptive plasticity in energetically costly stress responses (e.g., Brennan et al., 2022; Campbell-Staton et al., 2021). To differentiate between the gain of beneficial plasticity or canalization in benthic populations, future research could examine thermal responses of marine stickleback as a proxy for the ancestral state (Kirch et al., 2021). Indeed, recent work suggests that marine fish suffer negative effects from prolonged heat exposure (e.g., Spence-Jones et al., 2025). This study also provides evidence that transgenerational effects may ameliorate heat stress in stickleback, which we could not test in our experiment given that the parents were wild-caught. Additionally, the OCLTTH predicts that cool-adapted ectotherms evolve greater mitochondrial capacity to compensate for cold-induced reduction of ATP synthesis, leading to a higher SMR and lower CT_max_ (Pörtner, 2001). Indeed, we found that limnetic fish had a higher SMR at the secondary timepoint and a lower heat tolerance than benthic fish. However, the SMR of benthic and limnetic fish were nearly identical at the primary timepoint. These hypotheses could be tested by measuring temperature related shifts in energy reserves across the two ecotypes, incorporating a marine population as a proxy for the ancestral state, and through more detailed biochemical and transcriptomic analyses.

While our discussion has focused on heat itself as the primary selective agent, thermal tolerance variation may be correlated with evolution of other metabolically costly phenotypes. For example, previous work identified morphological and physiological changes connected to stickleback migration habits, with nonmigratory fish evolving lower MMR than migratory ecotypes (Dalziel et al, 2012a; Dalziel et al., 2012b). Similarly, migration-associated ecotype differences in SMR are also connected to heritable changes in thyroid hormone signaling (Kitano et al., 2010). Although all the lake fish in the present study are non-migratory, extending our results to a more geographically diverse set of populations could clarify the independent and combined effects of thermal environment and migration habits on metabolic evolution.

### Temperature did not affect infection rates or fibrosis despite metabolic costs of immunity

Temperature impacts immune responses across a wide range of teleosts (Hao et al., 2024; Scharsack & Franke, 2022), including stickleback (Dittmar et al., 2014; Franke et al., 2019; Scharsack et al., 2021). Thus, we were surprised to find no connection between temperature and infection or fibrosis. This negative result may be partly due to our recovery of only a single infected fish. Our experiment aimed to limit the influence of local coevolution between fish and parasites by using a foreign cestode (i.e., from Walby Lake), but this decision may explain the near total lack of infection in our study (i.e., stickleback tend to be better at resisting infection from foreign populations of *S. solidus* (Bolnick et al., 2024; Weber et al., 2017a)). However, fibrosis was associated with lower MMR, SMR, and AS at the end of the experiment (Figure 3). Fibrosis is initiated within days of initial cestode exposure (Hund et al., 2022) but can have long-term effects on mobility (Matthews et al., 2023). One possible interpretation is that lower AS results from oxygen deprivation due to reduced mobility or other metabolic costs of fibrosis.

### Partial support for the OCLTTH

In support of the OCLTTH, we found that AS declined in limnetic populations at higher temperatures. However, AS peaked at 22°C in limnetic fish at the start of the experiment, suggesting that 22°C is the T_opt_ for limnetic populations. This trend agrees with work showing that stickleback AS increases up to at least 20°C (Ressel et al., 2021). Yet, both the historical lake temperature and developmental temperature of these populations is well below 22°C, and previous work in other stickleback ecotypes suggests their T_opt_ is between 13-17°C (Barrett et al., 2011; Dittmar et al., 2014; Franke et al., 2017, 2019). In contrast to the OCLTTH, this implies that AS-based estimates of T_opt_ are not closely connected to environmental temperatures. A final disconnect with OCLTTH involves the expectation of correlated shifts between AS and body condition. Body condition and hepatosomatic index were the only two condition metrics associated with AS. Additionally, the expected positive association with AS was only present in benthic fish at the end of the experiment, while limnetic populations exhibited a negative trend for both of these condition metrics. Although the inconsistent relationship between AS and condition suggests that the OCLTTH may have limited relevance for stickleback, another explanation involves differences in energetic demand and uptake between lab and wild environments. Costs may be masked when resources are plentiful but apparent when they are scarce, as is likely the case in natural settings (Boots, 2011). Our laboratory fish were fed to satiation, perhaps leading to higher MMR than would be seen in resource-limited animals. Additionally, metabolic rate is highly plastic and varies within an individual’s lifetime, with potentially different fitness effects at different ontogenic stages. Indeed, one experiment found that zebrafish in food-limited, warm conditions produced offspring with lower metabolic and growth rates – adaptive traits in a nutrient-limited environment (Pettersen et al., 2024). Further tests of the OCLTTH will benefit from testing the impact of resource limitation on performance across temperatures, dietary settings, and ontogenic stages.

### Limitations

Several features of our experimental design limited our ability to test connections between temperature and metabolism. First, we were unable to individually mark the fish used in this experiment, and therefore could not directly connect AS at the start of the experiment to survival, growth rate, or individual-specific AS at the end of the experiment. Second, our choice to acclimate fish to constant temperatures was practical but not ecologically realistic. The chronic impact of high temperatures in this experiment likely exceeds what stickleback experience in the wild. Conversely, acclimating fish to high temperatures also appears to buffer the detrimental impacts of thermal stress compared to rapid environmental change (Earhart et al., 2022). Rapid temperature fluctuations are expected to become more frequent, eliminating the opportunity to acclimate (Diffenbaugh & Field, 2013). Although technically challenging, future studies that mimic daily oscillations in temperature may provide more accurate predictions regarding impacts of natural heat waves (e.g., Ruiz-Raya et al., 2026).

Both MMR (Ressel et al., 2021) and SMR (Chabot et al., 2016) can be difficult to measure using standardized methods, which also compromises estimates of AS. Our MMR assays were limited to exercising fish for one minute in compliance with animal care and ethics guidelines. Ideally, fish will be unresponsive to agitation of the caudal fin before MMR is measured (Clark et al., 2013). Some, but not all, fish in this experiment reached this state of exhaustion. While we attempted to negate this effect by removing a few outliers with artificially low MMR or high SMR (see supplemental methods), our observations of increases in primary MMR with temperature could have resulted from fish not being reliably exercised to exhaustion. Despite this limitation, all fish were exercised for an equal amount of time and our 20°C MMR values were higher than those reported from wild stickleback assayed at the same temperature and collected from the same region of south central Alaska (Ressel et al., 2021), which suggests that we did thoroughly exercise these animals. In the case of SMR, we did not collect behavioral data during the course of the assays, and some animals may have failed to reach inactive states. This could explain why some treatments (e.g. 20°C SMR) deviate from linear expectations. Importantly, while our metabolic rate estimates may not be perfectly comparable with other studies, they were still comparable between treatment groups.

## Conclusions

In contrast to previous work on short term thermal tolerance, this study demonstrates that threespine stickleback display large differences in long term thermal tolerance. These differences are closely connected to ecotype variation in thermal habitats, suggesting that they may result from local adaptation. Our work also provides a valuable test of the OCLTTH and suggests that AS stability underpins resilience to heat in benthic stickleback. Additionally, we find a clear impact of immunity on metabolism, with cestode exposure driving a fibrosis response that significantly lowered respirometry rates regardless of ecotype. Future studies should aim to identify the mechanistic basis of AS stability and whether benthic and limnetic ecotypes commonly evolve differences in chronic thermal tolerance.

## Permits/Acknowledgments/Funding

All experiments followed the National Research Council’s Guide for the Care and Use of Laboratory Animals and were approved by the Institutional Animal Care and Use Committees of the University of Wisconsin – Madison (Protocol #L006460). B. Solin, M. Chotlos, J. Abels, M. Reynolds, C. Wolf, and J. Benavente assisted with animal care and dissections. KTS, EVK, and JNW were supported by the NIGMS of the NIH (#1R35GM142891-01 to JNW) and lab startup funding from the University of Wisconsin-Madison (to JNW). EVK was supported by the UW-Department of Integrative Biology Graduate Summer Research Award.

## Author Contributions

All authors contributed to the design of the study and writing of the manuscript. KTS and EVK led data collection and analysis. JNW assisted in analysis and supervised and funded the work.

