## Supplemental Information for "Metabolic stability underpins thermal tolerance in benthic ecotypes of threespine stickleback (*Gasterostues aculeatus)*"

### **Supplemental Methods:**

#### *Animal husbandry and thermal manipulations*

Experimental fish were generated via in vitro fertilization of eggs from wild parents in Spring 2022. Eggs were shipped to the University of Wisconsin-Madison where they were kept at air temperature submerged in methylene blue-treated reverse osmosis water until hatching. Hatched fry were housed with siblings in 23L tanks until the start of the experiment. Aquaria temperature prior to the experiment was maintained at 17 ± 1°C. We fed fry to satiation once per day with *Artemia* nauplii and adults with pelletized fish feed (GP500, Brine Shrimp Direct; Ogden, USA). A 16 hour light: 8 hour dark circadian cycle was maintained for the duration of the experiment.

We used three criteria to determine the experimental temperature range (18-26℃). First, the maximum temperature is below the reported CT_max_ for this species, with estimates differing based on the length of trials and the use of acclimation treatments. CT_max_ in acute trials (i.e., rapid increases in temperature until loss of mechanical equilibrium) ranges from 32-34℃ (Barrett et al., 2011; Dammarck et al, 2018; Metzger et al., 2017; Mottola et al., 2022), while three longer term studies (using similar acclimation treatments and water chemistry as the current experiment) found limited mortality at 25℃ (4-week trial: Metzger & Schulte, 2017), 28℃ (14-day trial: Dittmar et al, 2014), and 50% mortality at 27-28℃ (10-day trial: Jordan & Garside, 1972). Our experimental temperature range also encompassed the projected maximum air temperatures in the Kenai, AK area during the hottest month of the year in 2060 under a moderate emissions scenario (RCP 6.0; https://snap.uaf.edu/tools/community-charts).

#### *Respirometry*

After a 24 hour fasting period, we anesthetized fish to record mass and standard length. Individuals were allowed to regain equilibrium in a holding tank, then chased with a net for one minute and transferred to 330mL respirometry chambers submerged in a temperature-controlled stock tank (TECO 2000 chiller; Hygger tank heater). Oxygenated water was intermittently flushed into respirometry chambers using automated software for 24 hours (AquaResp3: Morozov et al. 2019). We used oxygen-sensitive REDFLASH contactless sensor spots connected to a Firesting O2 oxygen meter (PyroScience GmbH) to record the temperature corrected oxygen concentration (mg/L) in each chamber.

#### *Statistical analysis*

All statistical analyses were conducted in R v. 4.3.2 (R Core Team, 2023). We used the FishResp package in R (Svendsen et al., 2019) to estimate SMR by first subtracting background microbial respiration, then averaging the lowest 20% of points from the entire data set after excluding outliers (i.e., a quantile approach (Chabot et al., 2016; Clark et al., 2013)). MMR was defined as the maximum value recorded during the trial (Clark et al. 2013), and AS is the difference between SMR and MMR. We then removed samples in which there were clear equipment malfunctions (i.e. negative SMR or MMR) or that died during the respirometry measurements (Supplemental Table 1). We also removed SMR and MMR outliers by calculating the mean and standard deviation of SMR/MMR across all temperature treatments and time points within each population. To remove individuals that were artificially stressed, SMR outliers were defined as ≥2SD above the mean. Meanwhile, MMR outliers were defined as ≥2SD below the mean to account for fish that were not exercised to exhaustion. If multiple fish from the same temperature treatment were flagged as outliers, they were retained to prevent erroneously eliminating temperature effects from the data. Using this method, three fish (two benthic, one limnetic) were removed from subsequent analysis.

Respirometry measurements were subset for each ecotype*temperature combination. Due to the quadratic nature of thermal performance curves (Clark et al., 2013; Pörtner et al., 2017), and because the thermal optima (i.e. the inflection point of a thermal performance curve) of experimental populations was unknown, we used AIC scores to compare linear models to models that included temperature as a quadratic term. We report quadratic results when the ΔAIC > 2 relative to the linear model. We used this same procedure to compare models that used mean depth, maximum depth, and surface area (in place of ecotype) as continuous predictors of MMR, SMR, and AS. We used binomial regression to test the effect of temperature and ecotype on fibrosis presence and ordinal regression (p-values based on 10,000 Monte-Carlo test permutations) to test the effect of temperature and ecotype on fibrosis severity (R package pgirmess v. 2.0.2; Giraudoux, 2024).

Allometry can severely bias estimates of body condition when there is significant between-group variation (Reist, 1986). While Fulton’s K remains a popular metric to assess body condition in stickleback, classification along the benthic-limnetic spectrum is often defined based on morphological variation, potentially biasing this condition metric from the outset. Therefore, in addition to using Fulton’s K, we also assessed body condition using scaled mass index (SMI; Pieg & Green, 2009). We calculated SMI both within ecotype and within populations (https://gist.github.com/chenpanliao/8ef652404d024c2c4191b8f159400a37#file-robust-scaledmassindex-r). Standard length was measured using digital calipers for all samples except for two Spirit Lake fish at the start of the experiment due to equipment failure.

### **Supplemental Tables:**

#### **Supplemental Table 1:** Sample sizes of fish used to measure SMR, MMR, and AS at the primary and secondary timepoints. Numbers in parentheses indicate the number of trials with equipment failure or mortality during the respirometry assay.

| **Ecotype** | **Population**  (Latitude, Longitude) | **Temperature (°C)** |  | **Primary Timepoint Sample Size** | **Secondary Timepoint Sample Size** |
| --- | --- | --- | --- | --- | --- |
| **Benthic** | **Watson**  (60.5365, -150.4628) | 18 |  | 16 | 15 (1) |
|  |  | 20 |  | 4 (8) | 0 |
|  |  | 22 |  | 15 (1) | 15 |
|  |  | 24 |  | 15 (1) | 0 |
|  |  | 26 |  | 13 | 11 |
|  | **Finger**  (61.6085, -149.2675) | 18 |  | 7 | 0 |
|  |  | 20 |  | 7 | 0 |
|  |  | 22 |  | 8 | 0 |
|  |  | 24 |  | 6 | 0 |
|  |  | 26 |  | 0 (6) | 0 |
| **Limnetic** | **Spirit**  (60.5951, -150.9995) | 18 |  | 15 | 15 |
|  |  | 20 |  | 15 | 0 |
|  |  | 22 |  | 14 (1) | 13 |
|  |  | 24 |  | 16 | 0 |
|  |  | 26 |  | 12 (4) | 0 |
|  | **Wik**  (60.7160, -151.2479) | 18 |  | 4 (1) | 0 |
|  |  | 20 |  | 7 | 0 |
|  |  | 22 |  | 7 | 0 |
|  |  | 24 |  | 7 (1) | 0 |
|  |  | 26 |  | 7 | 0 |

#### **Supplemental Table 2:** Model summaries for MMR, SMR, and AS at the primary (start) and secondary/tertiary (end) of the experiment.

| **Timepoint** | **Response** | **Predictors** | **AIC** | **R^2^_adj_** | **F** | **DF** | **p-value** |
| --- | --- | --- | --- | --- | --- | --- | --- |
| Primary | MMR ~ | Temperature * Ecotype | 2692.5 | 0.2462 | 20.82 | (3, 179) | <0.0001 |
|  |  | Temperature * Surface Area | 2690.669 | 0.2537 | 21.63 | (3, 179) | <0.0001 |
|  | SMR ~ | Temperature^2^ + Temperature + Ecotype | 2445.096 | 0.5788 | 84.37 | (3, 179) | <0.0001 |
|  |  | Temperature^2^ * Mean Depth + Temperature * Mean Depth | 2438.554 | 0.5979 | 55.12 | (5, 177) | <0.0001 |
|  | AS ~ | Temperature * Ecotype | 2653.389 | 0.01994 | 2.234 | (3, 179) | 0.08587 |
|  |  | Temperature^2^ * Surface Area + Temperature * Surface Area | 2634.064 | 0.1275 | 6.318 | (5, 177) | <0.0001 |
| Secondary/tertiary | MMR ~ | Temperature^2^ + Temperature + Ecotype + Sex + Fibrosis Presence | 1033.849 | 0.2847 | 6.413 | (5, 63) | <0.0001 |
|  |  | Temperature^2^ + Temperature + Ecotype + Sex + Fibrosis Score | 1037.56 | 0.2452 | 5.418 | (5, 63) | 0.0003305 |
|  | SMR ~ | Temperature^2^ + Temperature + Ecotype + Sex + Fibrosis Presence | 895.0609 | 0.475 | 13.31 | (5, 63) | <0.0001 |
|  |  | Temperature^2^ + Temperature + Ecotype + Sex + Fibrosis Score | 896.1145 | 0.4669 | 12.91 | (5, 63) | <0.0001 |
|  | AS ~ | Temperature^2^ + Temperature + Ecotype + Fibrosis Presence | 998.106 | 0.2845 | 7.76 | (4, 64) | <0.0001 |
|  |  | Temperature^2^ + Temperature + Ecotype + Fibrosis Score | 1001.381 | 0.2497 | 6.659 | (4, 64) | 0.0001518 |

#### **Supplemental Table 3:** Primary response of MMR, SMR, and AS were modeled using generalized linear models. Significant predictors (p<0.05) are bolded.

| **Response** | **Predictors** | **Estimate** | **SE** | **T value** | **P value** |
| --- | --- | --- | --- | --- | --- |
| **MMR** | Intercept | 3.691 | 310.569 | 0.012 | 0.991 |
|  | **Temperature** | **84.650** | **14.132** | **5.990** | **<0.0001** |
|  | Ecotype | 374.591 | 444.212 | 0.843 | 0.400 |
|  | Temperature*Ecotype | -13.273 | 20.236 | -0.656 | 0.513 |
| **SMR** | **Intercept** | **2567.493** | **1007.488** | **2.548** | **0.01166** |
|  | **Temperature** | **-246.710** | **93.596** | **-2.636** | **0.00913** |
|  | **Temperature^2^** | **7.494** | **2.145** | **3.494** | **0.00060** |
|  | Ecotype | 23.910 | 28.137 | 0.850 | 0.39658 |
| **AS** | **Intercept** | **653.27** | **279.09** | **2.341** | **0.0203** |
|  | Temperature | 17.70 | 12.70 | 1.394 | 0.1650 |
|  | **Ecotype** | **932.27** | **399.19** | **2.335** | **0.0206** |
|  | **Temperature*Ecotype** | **-39.65** | **18.19** | **-2.180** | **0.0305** |

#### **Supplemental Table 4:** Physical lake characteristics as predictors of respirometry traits at the start of the experiment. Notably, the depth data begins at zero and becomes more negative as a lake gets deeper. Significant predictors (p<0.05) are bolded.

| **Response** | **Predictor** | **Estimate** | **SE** | **T value** | **P value** |
| --- | --- | --- | --- | --- | --- |
| **MMR** | Intercept | -1444.027 | 906.019 | -1.594 | 0.112742 |
|  | **Temperature** | **157.224** | **41.144** | **3.821** | **^0.000183^** |
|  | Surface Area | 170.203 | 88.861 | 1.915 | 0.057037 |
|  | **Temperature*Surface Area** | **-8.267** | **4.086** | **-2.023** | **0.044547** |
| **SMR** | **Intercept** | **5508.9948** | **2107.0577** | **2.615** | **0.00970** |
|  | **Temperature** | **-495.3290** | **195.3893** | **-2.535** | **0.01211** |
|  | **Temperature^2^** | **12.5867** | **4.4648** | **2.819** | **0.00536** |
|  | Mean Depth | 426.1368 | 264.8567 | 1.609 | 0.10941 |
|  | Temperature * Mean Depth | -36.1191 | 24.4262 | -1.479 | 0.14100 |
|  | Temperature^2^ * Mean Depth | 0.7394 | 0.5558 | 1.330 | 0.18512 |
| **AS** | **Intercept** | **-29154.358** | **6941.585** | **-4.200** | **p<0.0001** |
|  | **Temperature** | **2744.643** | **646.506** | **4.245** | **p<0.0001** |
|  | **Temperature^2^** | **-60.905** | **14.869** | **-4.096** | **p<0.0001** |
|  | **Surface Area** | **2594.858** | **693.715** | **3.741** | **0.000248** |
|  | **Temperature*Surface Area** | **-234.376** | **64.948** | **-3.609** | **0.000400** |
|  | **Temperature^2^ * Surface Area** | **5.167** | **1.503** | **3.439** | **0.000729** |

#### **Supplemental Table 5:** Secondary/tertiary response of MMR, SMR, and AS were modeled using generalized linear models. Significant predictors (p<0.05) are bolded.

| **Response** | **Predictor** | **Estimate** | **SE** | **T value** | **P value** |
| --- | --- | --- | --- | --- | --- |
| **MMR** | **Intercept** | **15246.772** | **3088.403** | **4.937** | **p<0.0001** |
|  | **Temperature** | **-1305.376** | **292.022** | **-4.470** | **p<0.0001** |
|  | **Temperature^2^** | **30.419** | **6.785** | **4.483** | **p<0.0001** |
|  | **Ecotype** | **356.246** | **110.204** | **3.233** | **0.00195** |
|  | Sex | 63.501 | 100.217 | 0.634 | 0.52862 |
|  | **Fibrosis Presence** | **-324.169** | **108.315** | **-2.993** | **0.00394** |
|  | **Intercept** | **14371.697** | **3182.580** | **4.516** | **p<0.0001** |
|  | **Temperature** | **-1226.065** | **301.022** | **-4.073** | **0.000132** |
|  | **Temperature^2^** | **28.604** | **6.991** | **4.091** | **0.000124** |
|  | **Ecotype** | **336.360** | **112.736** | **2.984** | **0.004049** |
|  | Sex | 63.818 | 103.095 | 0.619 | 0.538136 |
|  | **Fibrosis Score** | **-134.977** | **59.248** | **-2.278** | **0.026117** |
| **SMR** | **Intercept** | **5892.511** | **1129.686** | **5.216** | **p<0.0001** |
|  | **Temperature** | **-527.506** | **106.817** | **-4.938** | **p<0.0001** |
|  | **Temperature^2^** | **13.040** | **2.482** | **5.254** | **p<0.0001** |
|  | Ecotype | 75.230 | 40.311 | 1.866 | 0.06666 |
|  | Sex | -55.269 | 36.658 | -1.508 | 0.13663 |
|  | **Fibrosis Presence** | **-116.943** | **39.620** | **-2.952** | **0.00444** |
|  | **Intercept** | **5531.523** | **1141.939** | **4.844** | **p<0.0001** |
|  | **Temperature** | **-494.506** | **108.010** | **-4.578** | **p<0.001** |
|  | Temperature^2^ | 12.288 | 2.509 | 4.898 | 0.7265 |
|  | **Ecotype** | **71.367** | **40.451** | **1.764** | **0.0099** |
|  | Sex | -53.698 | 36.992 | -1.452 | 0.6725 |
|  | **Fibrosis Score** | **-58.704** | **21.259** | **-2.761** | **0.0071** |
| **AS** | **Intercept** | **9391.38** | **2399.38** | **3.914** | **0.000223** |
|  | **Temperature** | **-772.61** | **226.86** | **-3.406** | **0.001144** |
|  | **Temperature^2^** | **17.20** | **5.27** | **3.263** | **0.001770** |
|  | **Ecotype** | **271.49** | **85.39** | **3.179** | **0.002276** |
|  | **Fibrosis Presence** | **-198.96** | **83.98** | **-2.369** | **0.020860** |
|  | **Intercept** | **8909.006** | **2464.451** | **3.615** | **0.000592** |
|  | **Temperature** | **-729.349** | **233.135** | **-3.128** | **0.002645** |
|  | **Temperature^2^** | **16.205** | **5.414** | **2.993** | **0.003922** |
|  | **Ecotype** | **255.347** | **87.067** | **2.933** | **0.004657** |
|  | Fibrosis Score | -70.626 | 45.726 | -1.545 | 0.1284 |

### **Supplemental Figures:**

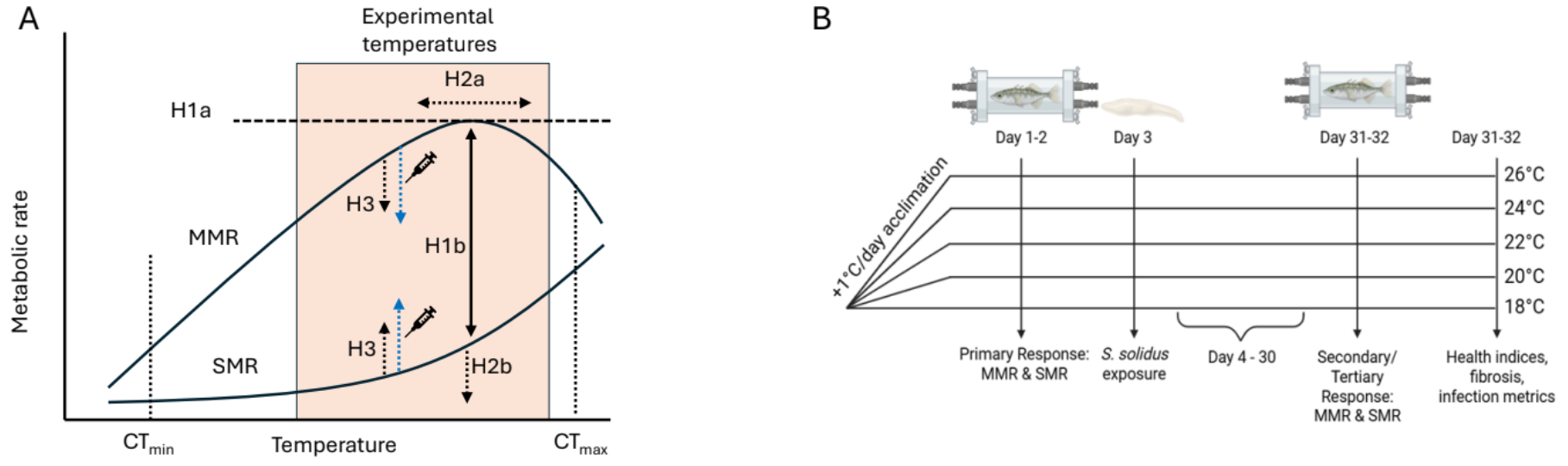

##### **Supplemental Figure 1. (A)** Metabolic rates change with temperature to meet respiratory demands, and the critical thermal limits (CT_min_ and CT_max_) reflect boundaries where there is sufficient energy to maintain homeostasis. The OCLTTH predicts that maximum metabolic rate (MMR) cannot continually increase with temperature (H1a), leading to a temperature at which aerobic scope is maximized (H1b: AS_max_). By exposing stickleback to temperatures near their predicted MMR inflection point (orange box), but well below CT_max_, we expected that fish from thermally dynamic lakes would display greater thermal breadth (i.e., stable AS across a range of high temperatures) and more stable body condition at high temperature. AS stability could be achieved by either broadening the peak of their MMR curve (H2a) or increasing SMR more slowly with temperature (H2b). If immune responses or parasite infections require greater investment in SMR or reduce MMR , then these should also alter thermal performance by decreasing AS and body condition in an ecotype-specific manner (H3, blue arrows represent limnetic fish, black arrows are benthic fish, fibrosis is represented by the syringe). **(B)** Summary of experimental design (graphic created on BioRender).

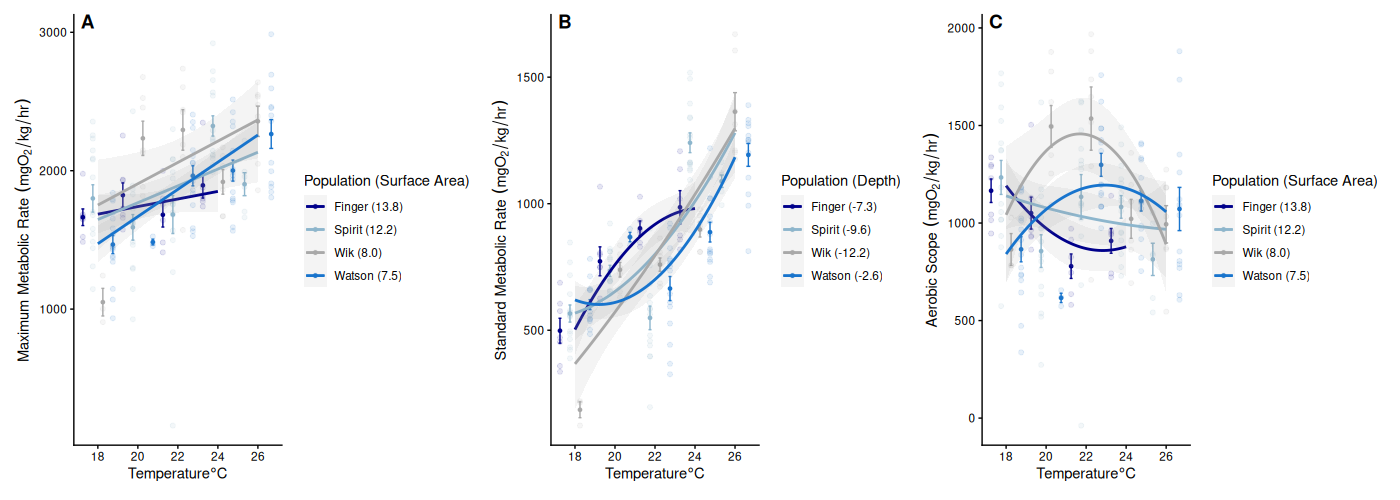

#### **Supplemental Figure 2:** Primary measurements of **(A)** MMR, **(B)** SMR, **(C)** and AS as a function of temperature for each lake. MMR and AS were best predicted by lake surface area (ha). SMR was best predicted by mean lake depth (m). Depth begins at 0m and becomes more negative with a deeper lake.

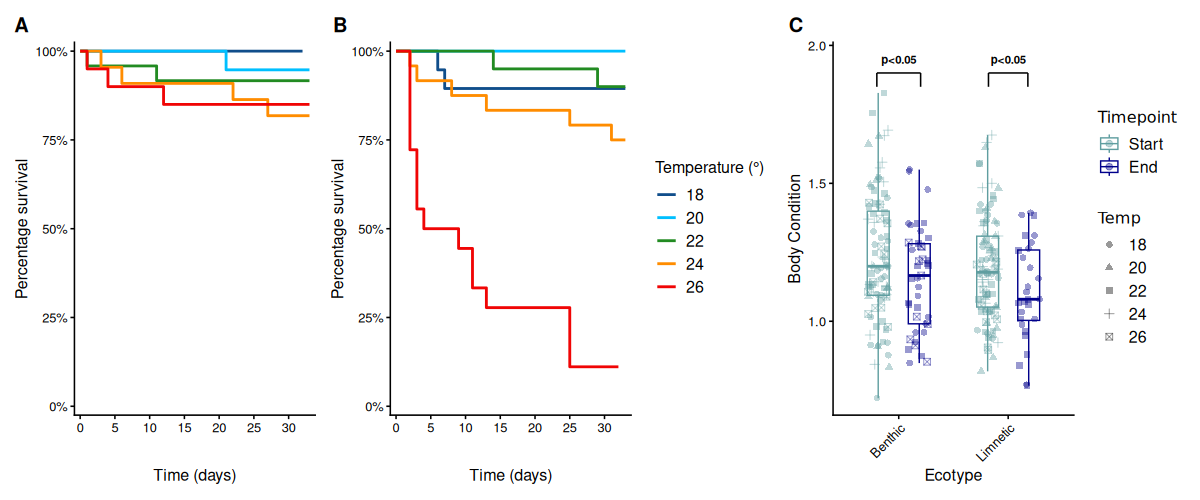

**Supplemental Figure 3:** Declines in body condition between the start and end of the experiment for benthic and limnetic fish.

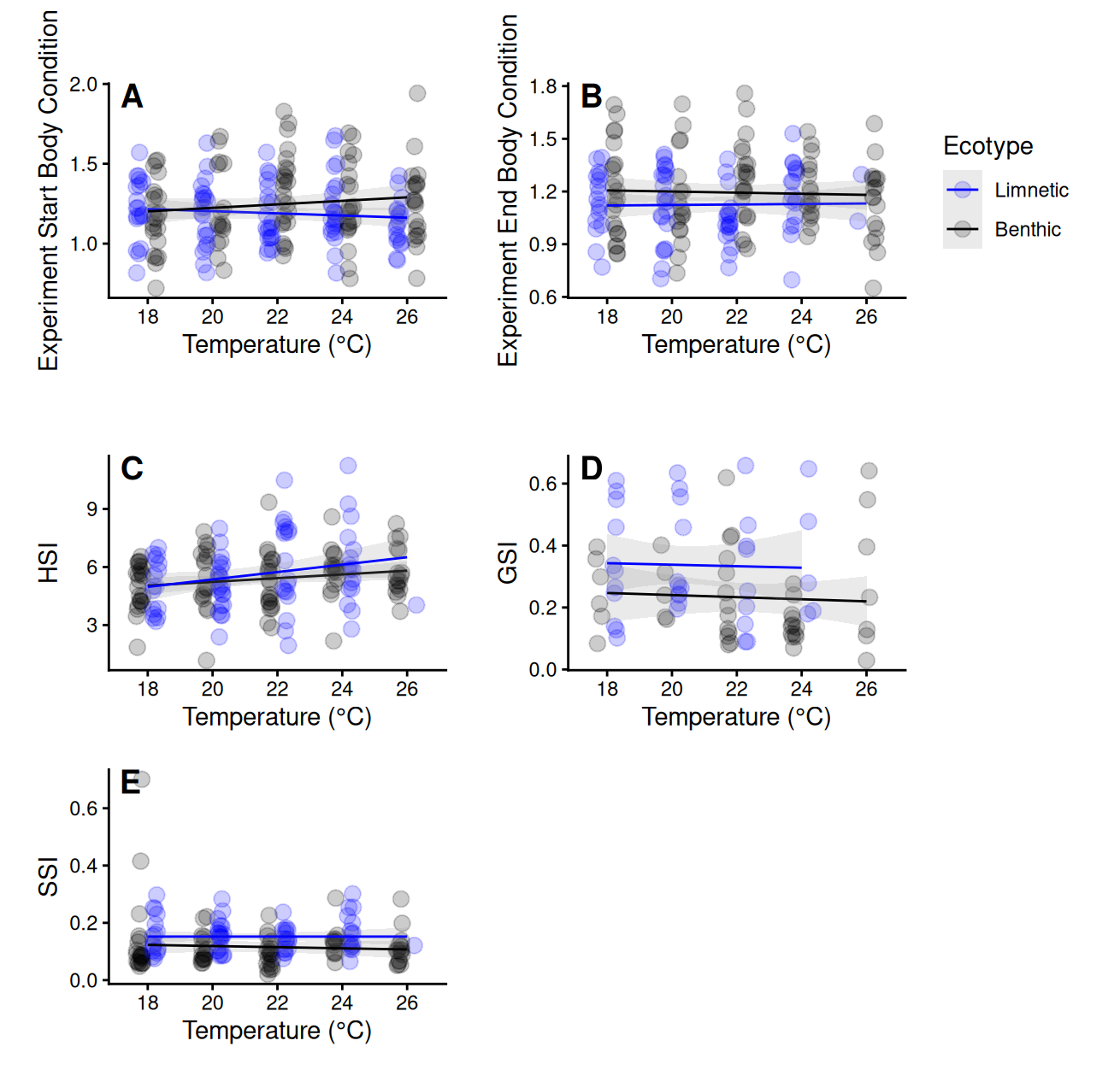

#### **Supplemental Figure 4:** Metrics of condition as a function of temperature for each ecotype. **(B)** Body condition at the secondary time point decreased slightly more in benthic fish than limnetic fish across temperatures. **(A)** Body condition at the primary timepoint, **(C)** HSI, **(D)** GSI, and **(E)** SSI had no association with temperature.

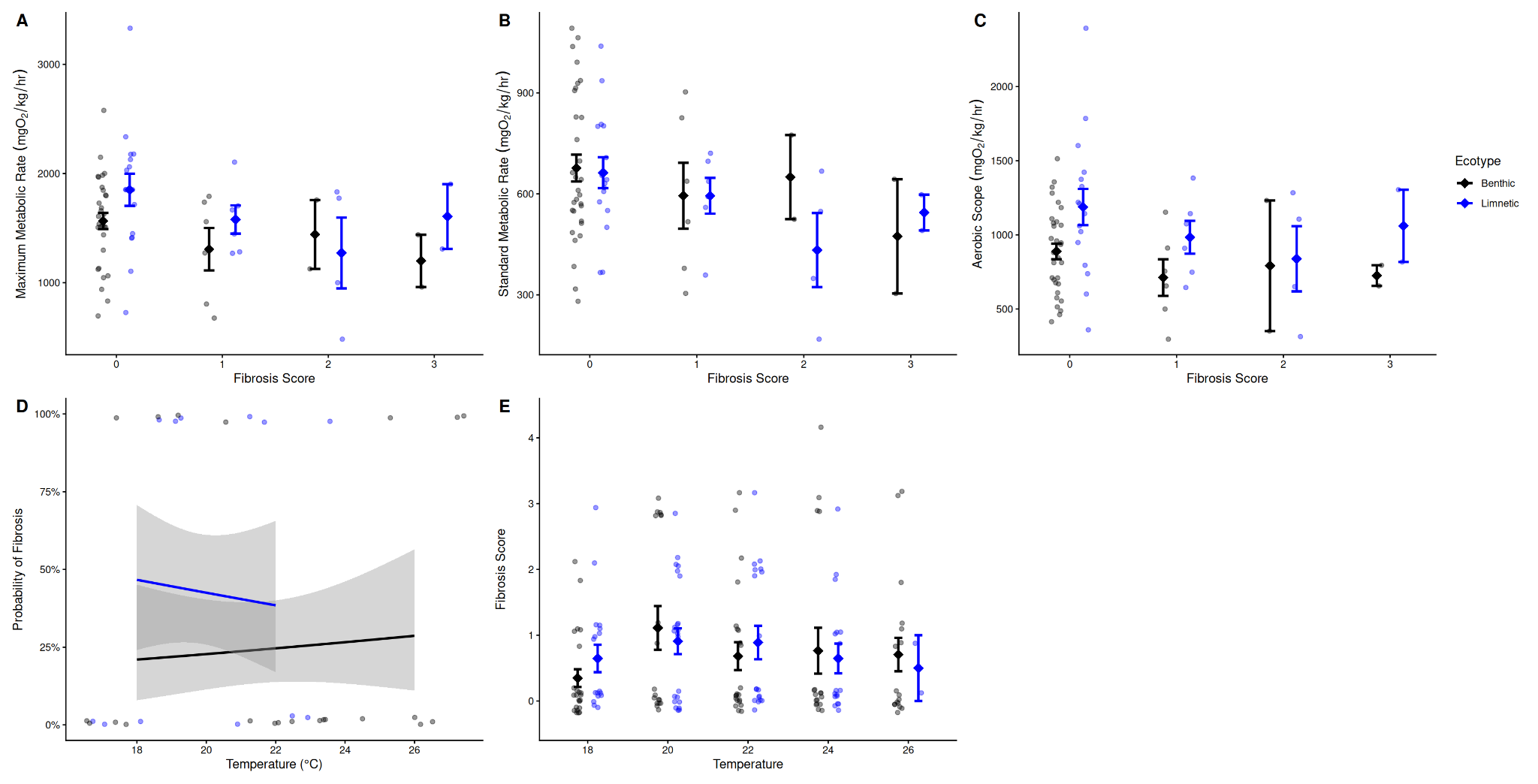

#### **Supplemental Figure 5:** **(A)** MMR, **(B)** SMR, and **(C)** AS as a function of fibrosis score. Fibrosis severity significantly decreased MMR (t = -2.278, p = 0.0256) and SMR (t = -2.761, p = 0.0071), but not AS. **(D)** Fibrosis presence and **(E)** severity were not influenced by temperature.

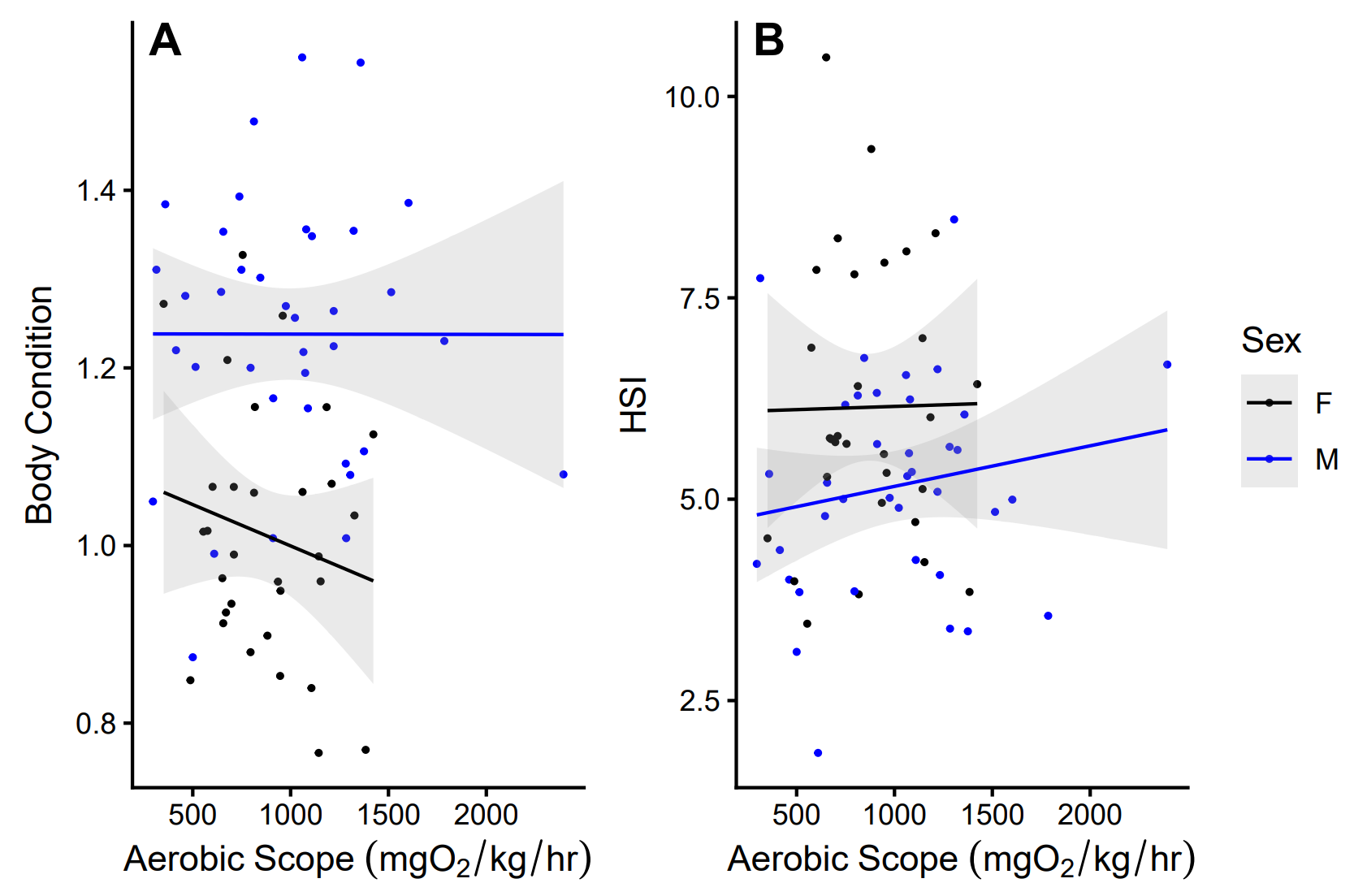

#### **Supplemental Figure 6: (A)** Fulton’s K body condition tended to be higher in males than females across AS (t = 5.971934, df = 61, p < 0.0001), while **(B)** HSI tended to be higher in females than males (t = -2.999606, df = 60, p = 0.0039).

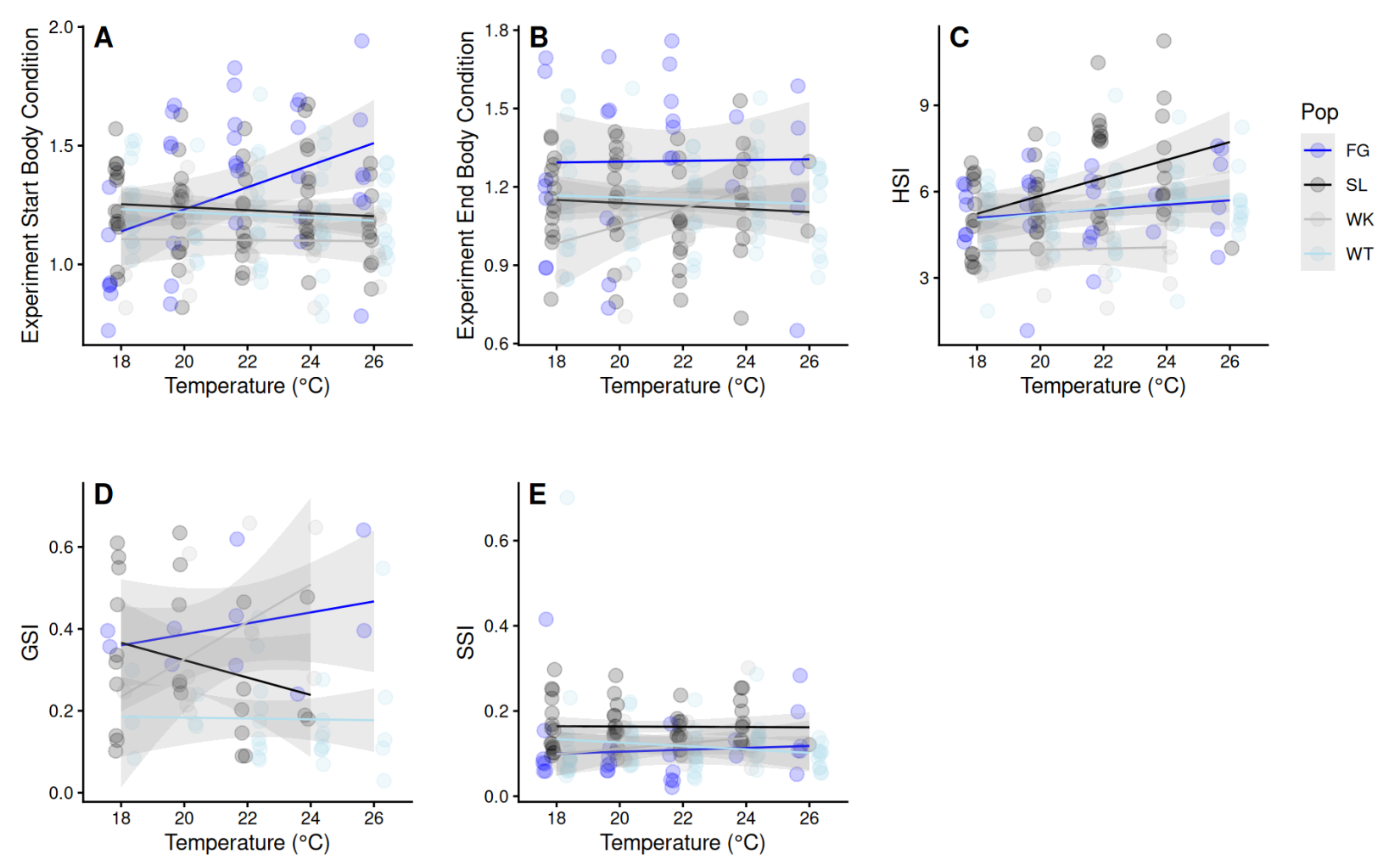

#### **Supplemental Figure 7:** The effect of temperature on condition metrics by population. **(A)** Finger Lake had a significantly different association between temperature and Fulton’s K at the start of the experiment than all other populations (p_adj_ < 0.05 for all post hoc slope pairwise comparisons between populations), **(B)** but this difference disappeared by the end of the experiment. There was no population-specific effect of temperature on **(C)** HSI, **(D)** GSI, or **(E)** SSI.

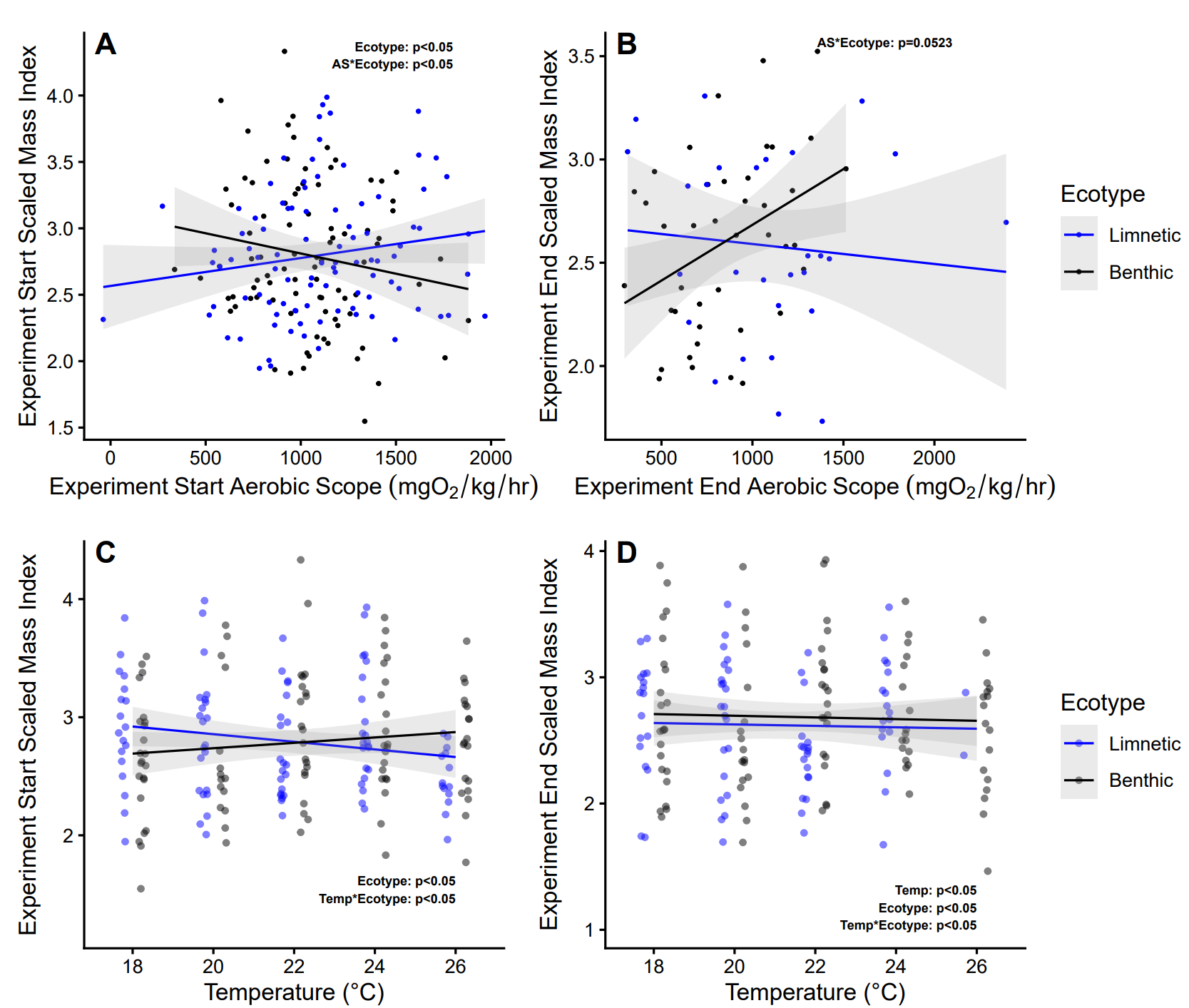

#### **Supplemental Figure 8:** The relationship between SMI and AS or temperature. **(A)** As with Fulton’s K, SMI increased with AS in limnetic fish but decreased among benthic fish at the primary timepoint, **(B)** while this trend flipped by the end of the experiment. **(C)** There was a significant temperature*ecotype interaction associated with temperature at the start of the experiment, with benthic fish increasing SMI with heat while limnetic fish displayed a decrease in SMI. **(D)** However, by the end of the experiment both ecotypes showed a slight but significant decrease in SMI with temperature, with the effect being greater in benthic fish.
